# N-WASP regulates the mobility of the B cell receptor and co-receptors during signaling activation

**DOI:** 10.1101/619627

**Authors:** Ivan Rey-Suarez, Brittany Wheatley, Peter Koo, Zhou Shu, Simon Mochrie, Wenxia Song, Hari Shroff, Arpita Upadhyaya

## Abstract

Regulation of membrane receptor mobility is important in tuning the cell’s response to external signals. This is particularly relevant in the context of immune receptor signaling. The binding of B cell receptors (BCR) to antigen induces B cell receptor activation. While actin dynamics and BCR signaling are known to be linked, the role of actin dynamics in modulating receptor mobility is not well understood. Here, we use single molecule imaging to examine BCR movement during signaling activation and examine the role of actin dynamics on BCR mobility. We use a novel machine learning based method to classify BCR trajectories into distinct diffusive states and show that the actin regulatory protein N-WASP regulates receptor mobility. Constitutive loss or acute inhibition of N-WASP, which is associated with enhanced signaling, leads to a predominance of BCR trajectories with lower diffusivity and is correlated with a decrease in actin dynamics. Furthermore, loss of N-WASP reduces diffusivity of CD19, a stimulatory co-receptor of the BCR but not that of unstimulated FcγRIIB, an inhibitory co-receptor. The effect of N-WASP is mirrored by inhibition of the Arp2/3 complex and formins. Our results implicate the dynamic actin network in fine-tuning receptor mobility and receptor-ligand interactions, thereby modulating B cell signaling.

---

B cells are an important component of the adaptive immune system. B cells sense antigen using specialized receptors known as B cell receptors (BCRs) that trigger signaling cascades and actin remodeling upon binding antigen on the surface of antigen presenting cells (APC)^1–3^. Signaling activation results in spreading of the B cell on the APC surface leading to the formation of a contact zone known as the immunological synapse^4^.

Antigen crosslinking aggregates BCRs in lipid rafts, enabling lipid raft-resident Src kinase to phosphorylate their immunoreceptor tyrosine-based activation motifs (ITAMs)^5–7^. Signaling BCRs assemble into microclusters, which grow via movement of BCRs and their incorporation into these microclusters^1,8^. BCR clustering is dependent on the probability of receptor-receptor interactions at the plasma membrane^9^, and is in part dictated by the lateral mobility of receptors. Thus, elucidating the mechanisms that regulate BCR movement in the cell membrane is critical for understanding BCR signaling.

The cortical actin network in cells is known to form juxtamembrane compartments that can transiently confine the lateral movement of membrane proteins^10–12^, including BCRs in B cells^13^. Treanor et al.^13^ showed that in un-stimulated B cells, inhibition of actin polymerization leads to an increase in lateral diffusivity of BCR and is accompanied by signaling that is reminiscent of activation. The transient dephosphorylation of ezrin and actin depolymerization induced by BCR-antigen interaction results in the detachment of the cortical actin from the plasma membrane concurrent with a transient increase in the lateral movement of surface BCRs^14^. Activation of Toll-like receptors sensitizes BCR signaling, by increasing BCR diffusivity through the remodeling of actin by cofilin, an actin binding protein that disassembles actin filaments^15^. The sub-membrane actin cytoskeleton also modulates the concentration of inhibitory co-receptors^16,17^ in the vicinity of BCR microclusters, thereby ensuring the rapid inhibition of activated BCRs.

A consensus picture that emerges from these studies is that in resting B cells, the actin network serves as a structural barrier for BCRs, regulating their mobility by steric interactions. However, considerable evidence points to a role for actin dynamics in regulating BCR signaling and activation. A study by Liu et al. found that N-WASP (Neural Wiskott-Aldrich syndrome protein), which activates the Arp2/3 complex and promotes branched actin polymerization, plays an important role in the deactivation or attenuation of BCR signaling^18^. B cells from N-WASP conditional knockout mice exhibit delayed cell contraction and display higher levels of signaling for longer times than control cells. However, these studies have largely focused on changes in the dynamics of cell spreading and BCR microcluster coalescence on a global cell-wide scale. Whether and how actin dynamics directly modulate BCR diffusion and signaling is an open question^19^. Inhibition of actin polymerization by low concentrations of Latrunculin A following antigen stimulation inhibits the growth of BCR microclusters^20^, suggesting that actin dynamics plays a direct role in modulating BCR mobility. Given the role of N-WASP in modulating B cell signaling, we hypothesized that N-WASP mediated actin regulation will have a significant effect on BCR mobility.

Here, we used single molecule and super-resolution imaging to investigate the diffusive properties of the BCR, its stimulatory co-receptor, CD19, and its inhibitory co-receptor FcγRIIB^21,22^. To obtain a better understanding of the diffusive properties of these receptors, we used a statistical algorithm to classify single molecule trajectories into states with distinct diffusivities and correlate these with potential signaling states. Additionally, we used conditional N-WASP knockout (cNKO) B cells, which are known to exhibit enhanced activation, to investigate how altered actin dynamics manifest differences in BCR signaling and diffusivity. We found that BCR and CD19 molecules had an overall lower diffusivity in cNKO B cells than in control cells. This reduction in diffusivity was due to a change in the proportion of receptors with low mobility. However, FcγRIIB receptors did not show a difference in diffusivity or state distribution between control and cNKO cells, suggesting that the effect of N-WASP on BCR and CD19 mobility is not global for all membrane proteins during B cell activation. We found that inhibition of N-WASP also reduces BCR diffusivity and actin flows, suggesting that the effect of N-WASP on BCR mobility is actin mediated. Overall, our study reveals how actin dynamics regulate the nanoscale organization and diffusivity of BCR during activation.

## Results

### B cell receptor motion spans a wide range of diffusivity

Primary murine B cells were allowed to spread on a supported lipid bilayer coated with mono-biotinylated Fab’ fragment of BCR-specific antibody (mbFab) that induces BCR. We used Interference Reflection Microscopy (IRM) to visualize the spreading and contraction of B cells on supported lipid bilayers (Fig. 1a, top panels), and total internal reflection fluorescence (TIRF) microscopy to analyze the clustering of BCR and coalescence of BCR clusters during cell contraction (Fig. 1a, bottom panels). B cell receptor diffusivity was extracted from single molecule imaging of BCR trajectories obtained by TIRF imaging (Fig. 1b). We verified that B cells underwent signaling activation in our experimental conditions by labeling with phosphotyrosine as well by quantifying the intensity increase of BCR clusters due to coalescence (Supplementary Fig. S1). To label the BCR, Alexa Fluor (AF) 546 labeled mbFab was added to the imaging medium at low concentrations (<1 μM) so that only single molecules were detected^23^. Cells were imaged from the moment they contact the bilayer and time-lapse movies (1000 frames at 33 Hz) were recorded. Figure 1c shows a representative frame showing single BCR molecules and the cell outline obtained from an interference reflection microscopy (IRM) image taken immediately after the time-lapse movie. Single molecules detected in each frame are localized with high precision (~20 nm) and linked frame by frame to generate tracks^24^. Only single molecules identified within the IRM based cell contour are taken into consideration. A representative compilation of the tracks obtained from a single cell during a 10 minute period is shown in Figure 1d. The tracks are color coded for short-time diffusivity, calculated using the method presented in Flyvbjerg et al^25^. The cumulative distribution of diffusion coefficients measured across the population of tracks show variation in diffusivity over several orders of magnitude. These results indicate that BCR exhibit a wide spectrum of diffusivities, which may be indicative of their signaling properties and their biochemical state. Moreover, we found the there was a larger proportion of BCR with higher mobility in the first minute compared to later time points (Fig. 1e). This is further reflected in the box plot, which compares the diffusivity distributions measured at the indicated time points (Fig. 1f). The diffusivity at minute 1 was significantly higher than at subsequent time points.

**Figure 1.**
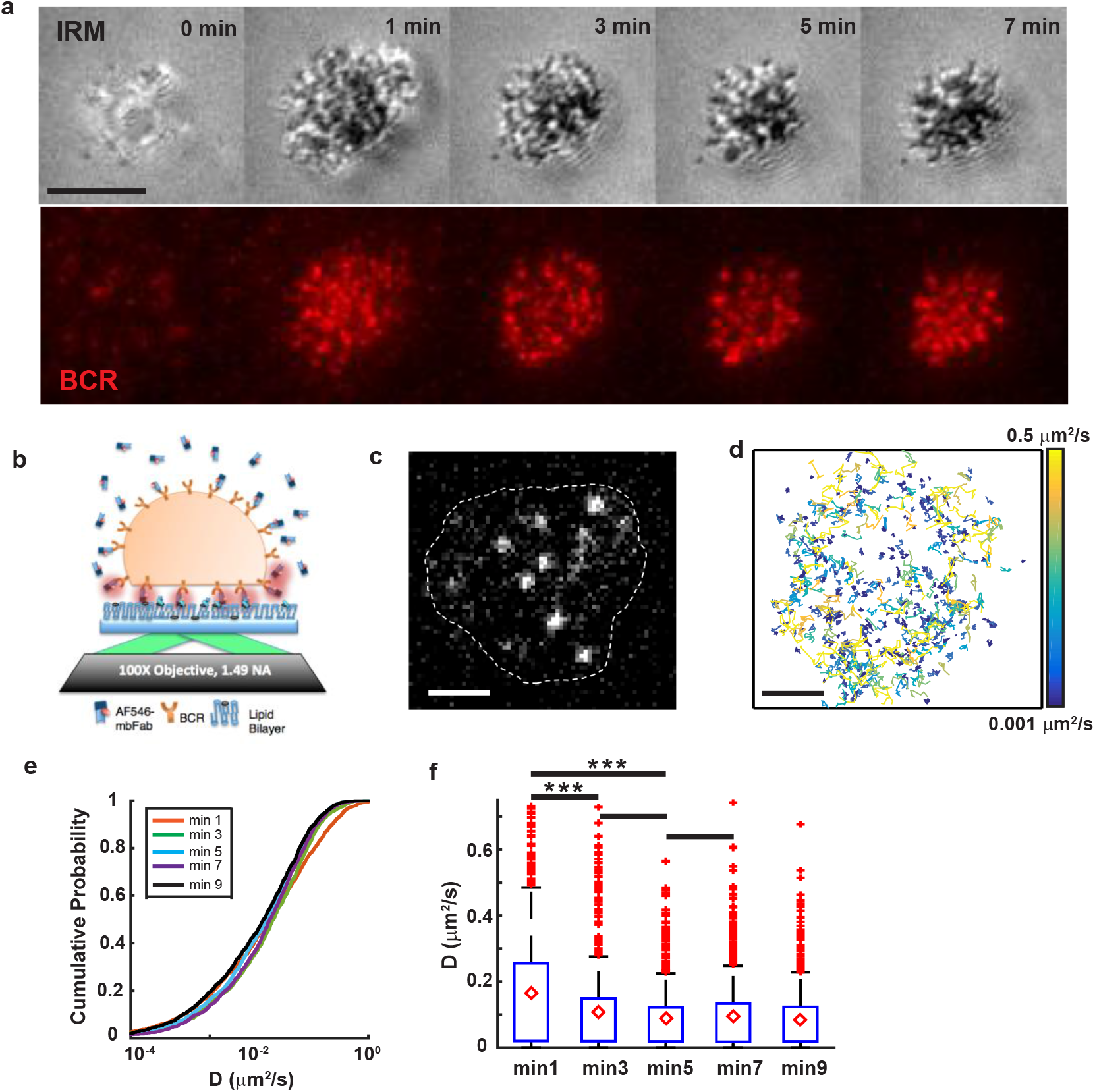
Single particle tracking reveals a wide range of BCR mobility. **a)** Panel showing primary murine B cell spreading with IRM (above) and BCR clustering (TIRF, below). Scale bar is 3 μm. **b)** Experimental schematic, indicating activated murine primary B cells, placed on supported lipid bilayers coated with mono-biotinylated fragments of antibody (mbFab). Cells are imaged in TIRF mode and the concentration of AF546 labeled mbFab is kept low enough to image single molecule events. **c)** Representative TIRF image with the bright dots representing single BCR molecules. The cell contour is obtained from an IRM image taken after TIRF imaging. Scale bar is 1 μm. **d)** The collection of tracks obtained for a control cell during a 10-minute period. The tracks are color-coded for diffusivity. Scale bar is 1 μm. **e)** Cumulative distribution function for the diffusivities measured at 1, 3, 5, 7 and 9 minutes after activation for BCR in B cells from control mice. **f)** Boxplot showing BCR diffusivities at the indicated time points where the mean is marked with red diamonds (N = 15 cells). Significance of differences was tested using the Kruskal-Wallis test (***P < 0.001; min 1 vs min 3, P = 0.0008; min 1 vs min 5, P = 0.000038; min 3 vs min 5, P = 0.1767; min 5 vs min 7, P = 0.8614).

### Perturbation-Expectation Maximization analysis identifies distinct diffusive states for BCR

In order to obtain a better understanding of the diffusive properties of the BCR, we employed a classification algorithm. Perturbation-Expectation Maximization (pEM) is a systems level analysis that uses machine learning algorithms to extract the set of distinct diffusive states that best describes a diffusivity distribution^26,27^. pEM works on the premise that for a protein, different biochemical interactions lead to different diffusive behaviors (diffusive states) and uses a classification strategy to uncover the different interactions in a statistically rigorous manner.

Here, we used pEM-v2, which is a pEM version capable of accounting both for non-normal diffusive modes and for the high heterogeneity of the cell membrane environment by splitting the trajectories into shorter segments and then identifying transitions between different diffusive states from one segment to the next. All the single molecule trajectories obtained were split into 15 frame segments and the classification analysis was performed on the set of all of these15-frame-long tracks. The trajectories of BCR from control cells were analyzed using pEM and 8 distinct states were identified. A number of representative trajectories assigned to each of these states are collected in Figure 2a in a 1.5 × 1.5 μm^2^ window.

**Figure 2.**
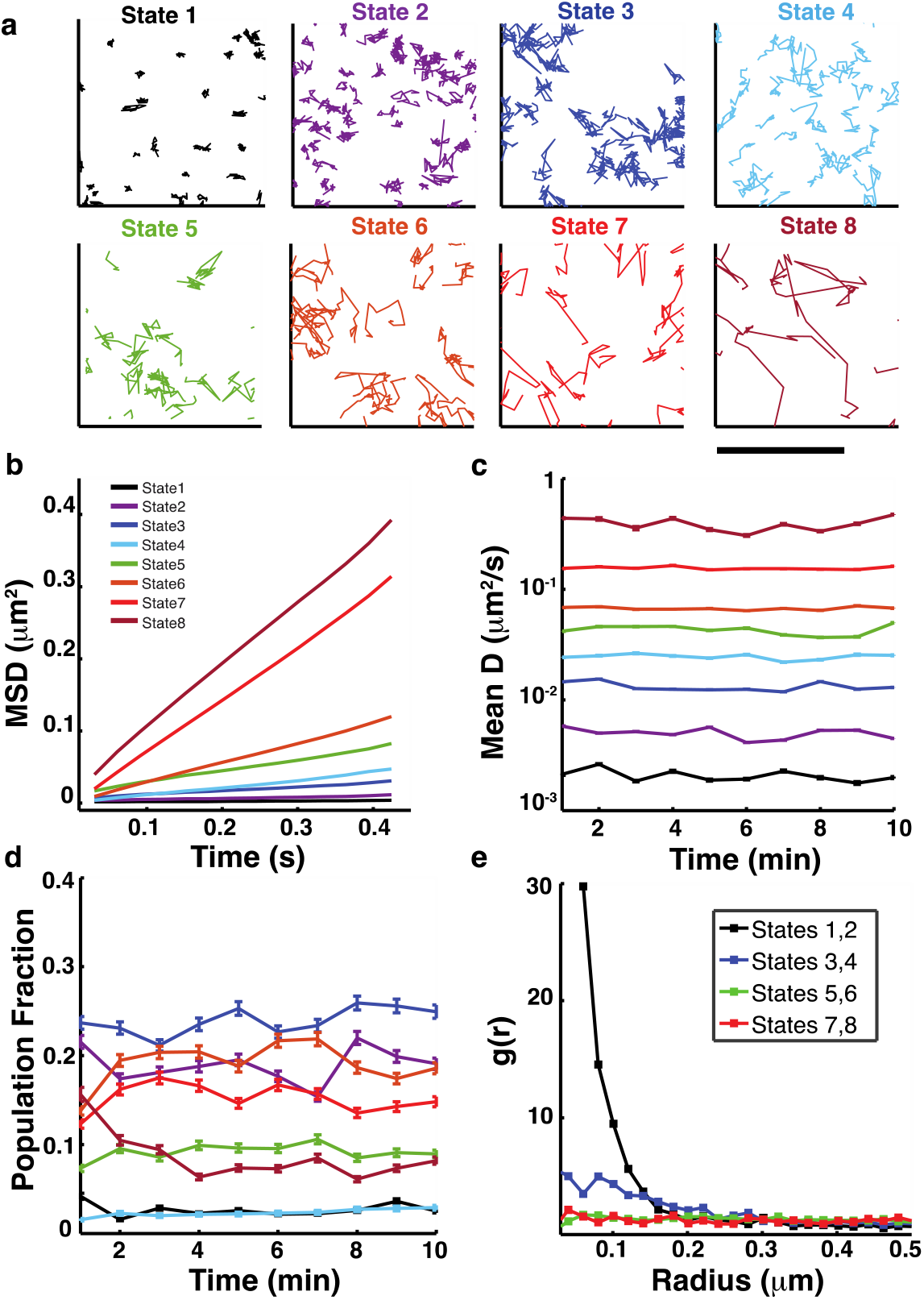
Perturbation Expectation Maximization analysis identifies eight distinct diffusive states for BCR in control cells. **a)** Characteristic tracks belonging to each of the BCR diffusive states identified by pEM. Diffusivity increases from state 1 (slowest) to state 8 (fastest). Scale bar is 1 μm. **b)** Ensemble mean square displacement (eMSD) plots for each of the states. Colors corresponding to different states are as shown in the legend. The same color scheme is used for all subsequent figures. (N = 15 cells). **c)** Plot showing the mean diffusivity for the trajectories belonging to each state at every time point. Error bars represent the standard error of the mean. **d)** Plots showing the fraction of BCR tracks that are sorted in each state at every time point. Error bars represent a confidence interval of 95% on the population fraction calculation. **e)** Plot of pair correlation as a function of distance for all states.

In control cells, all the 8 states displayed simple diffusion, as the ensemble mean square displacement (eMSD) plot of each state is linear (Fig. 2b). The mean diffusivity for all time points considered for each of these 8 states is shown in Figure 2c. For all states, the representative mean diffusivity is preserved over time. For each time point, we also calculated the fraction of tracks assigned to each state - the population fraction (Fig. 2d). The population fraction of the fastest moving state, state 8, rapidly reduces within the first few minutes and then stabilizes. This corresponds to the decrease in diffusivity for the higher mobility fraction as seen in the cumulative distribution function in Figure 1e. The population fraction corresponding to most of the other states fluctuate over time but do not show a clear trend. The first position of each non-redundant trajectory (for each distinct particle) was used to compute the spatial pair correlation function as a function of distance, *g(r)*, between the localized spots (Fig. 2e)^28^. *g(r)* measures the normalized probability of finding a second localized fluorophore (initial point of a trajectory) at a given distance, *r*, from an average localized fluorophore. For the resulting curves, a value of 1 indicates that the receptors are randomly organized, whereas values > 1 denote clustering. The range *r* over which *g(r) >1* denotes the scale of clustering. To calculate the pair correlation functions from our data, we combined the trajectories belonging to pairs of states that are closest in diffusivity (e.g. states 1 and 2; states 3 and 4 and so on). We see from Figure 2e that the lowest mobility states, states 1 and 2, display a *g(r)* that is significantly larger than 1 for small values of *r*, suggesting that these trajectories are significantly more densely clustered compared to those of the other states. States 3 and 4 show a very small degree of clustering, while the other higher mobility states display a largely homogeneous distribution. Of note, the slowest diffusive states, states 1 and 2, appear to be the ones that correspond to BCR in clusters.

### B cell receptor diffusivity is reduced in activated B cells from N-WASP-KO mice

While actin reorganization is known to be essential for signaling activation, the role of actin regulatory proteins in modulating the mobility of B cell receptors has not been studied and is thus less understood. In order to investigate how BCR diffusivity is modulated by actin regulatory proteins, we measured the diffusion of BCR molecules in B cells from mice where N-WASP is exclusively knocked out in B cells. We used the same single molecule imaging technique as described for control cells to obtain tracks of single BCR over time in cNKO cells (Fig. 3a). The different track colors correspond to tracks with different diffusivity and show a preponderance of tracks with lower diffusivity (blue) as compared to BCR tracks in control cells. However, the decrease in diffusivity for the high mobility tracks in the first few minutes that was observed for control cells, is not evident for BCR molecules in cNKO cells (Fig. 3b).

**Figure 3.**
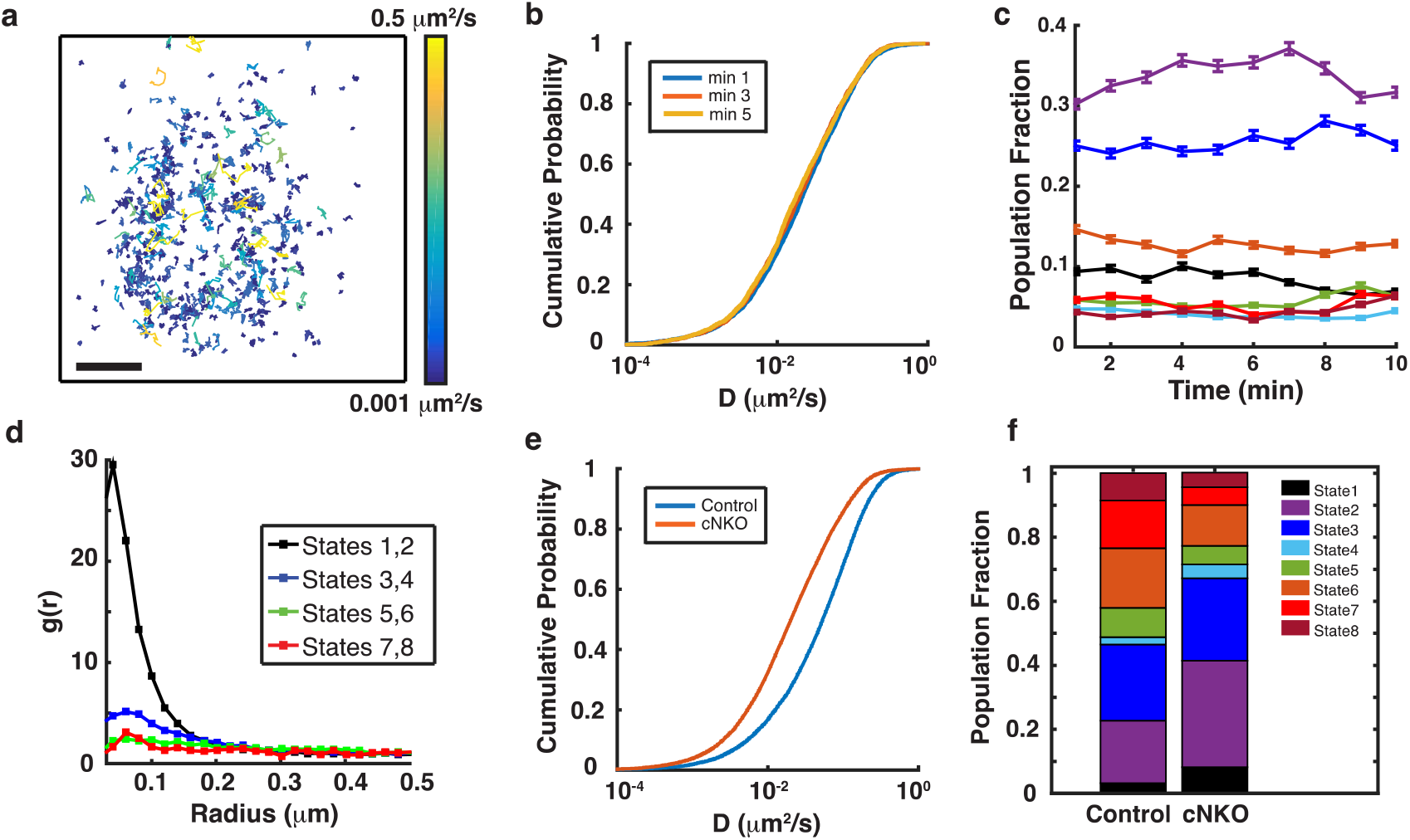
N-WASP knockout leads to predominance of BCR molecules in lower mobility diffusive states. **a)** Collection of tracks obtained from pEM analysis of BCR molecules in a cNKO cell during a 10 minute period. The tracks are color-coded for diffusivity. Scale bar is 1 μm. **b)** Cumulative distribution function for diffusivities measured at 1, 3 and 5 minutes after activation for BCR in B cells from cNKO mice. **c)** Plots of population fractions of eight distinct diffusive states as a function of time for BCR in cNKO cells. Error bars represent a confidence interval of 95% on the population fraction calculation. The colors corresponding to the different states are as shown in Figure 2b. **d)** Pair correlation function plots for different diffusive states. **e)** The distribution of diffusivities from the 5-10 minute time period after activation, for BCR in control and cNKO cells. The distributions for control and cNKO cells are significantly different (control cells N = 15, cNKO cells N = 17, P < 0.0001 Kruskal-Wallis test). **f)** Comparative population fractions for BCR in different states over the entire time period in control and cNKO cells. Significance values for the differences are in Supplementary Table 1.

To obtain more detailed insight into the differences between diffusivities in control and cNKO cells, we next used pEM analysis to assign BCR trajectories to diffusive states in cNKO B cells. The data containing the trajectories of BCR from control and cNKO cells was analyzed together and 8 different states were identified, as for the control case. Plots of the population fraction of BCR in cNKO cells as a function of time are shown in Figure 3c. The fraction of BCR in each state shows little change over time, and the eMSD of every state displays simple diffusion in cNKO B cells (Supplementary Fig. S2). Figure 3d shows the pair correlation function plots of all states in cNKO cells. Similar to control cells, states 1 and 2 display clustering, with states 3 and 4 showing a somewhat non-homogeneous distribution, while the higher mobility states have a homogenous distribution.

Given that the diffusivity remains largely unchanged for times beyond 5 minutes in both types of cells, we compared the cumulative distribution of diffusion coefficients measured for times between 5 and 10 minutes for control and cNKO cells. We found that BCR in cNKO cells display much slower diffusivity than control cells (Fig. 3e). This is also evident from the trajectories, which show a greater occurrence of tracks with lower diffusivity (blue color coded tracks, in Fig. 3a, compared to Fig. 1d). We found that, similar to control B cells, the lower mobility states, states 1-3 are the most dominant states in cNKO cells. We next compared the population fractions (pooled across time) of the occupied states for control and cNKO cells. We found significant differences (see Supplementary Table 1) in the population fractions between control and cNKO cells, with a significantly larger proportion of trajectories occupying the lowest mobility states (1-3) in cNKO cells (Fig. 3f).

In addition to identifying diffusive states, pEM analysis can also determine the transitions between these states along individual trajectories (Supplementary Fig. S3a). We thus selected longer tracks (>30 frames) and identified the state(s) to which the sub-tracks had been assigned and quantified the frequency of transitions from a given state to different states (Supplementary Fig. S3b, c). We found that BCRs tend to remain in their current states or switch to an adjacent one. BCR molecules in the three slowest diffusive states were the most stable in both control and cNKO cells, showing the least transition probability to other states. These observations suggest that fast diffusive particles will be more likely to encounter a cluster and be incorporated into it, thereby transitioning into the neighboring slower state. BCRs in cNKO cells showed an overall trend to transition towards slow diffusive states, especially towards states 2 and 3 (Supplementary Fig. S3c). In addition, particles in fast diffusive states were less stable and transitioned into slower states more frequently in cNKO than in control cells. This result is consistent with the higher population fraction of BCR in slow diffusive states (likely big clusters) in cNKO cells. Taken together, the predominance of lower mobility states of BCRs and the clustering of these low mobility states in cNKO cells which have been shown to have high levels of BCR signaling, are consistent with a model in which the lower diffusivity of BCR corresponds to signaling, clustering and activation.

### N-WASP modulates the diffusivity of the stimulatory co-receptor CD19 in activated cells

To better understand the nature of the BCR diffusive states that were enhanced in cNKO activated cells, we next investigated how the diffusivity of CD19, a stimulatory co-receptor^29^, is affected by the lack of N-WASP. CD19 is recruited to the BCR upon antigen binding, which enhances BCR activation. A previous study using super-resolution imaging found that in resting B cells, CD19 resides in nanoclusters separated from IgM BCR nanoclusters, while in activated B cells CD19 and BCR nanoclusters are colocalized^30,31^. Thus, single molecule studies of CD19 have the potential to reveal additional insight into signaling BCR states. Consistent with previous reports, CD19 and BCR microclusters co-localized to within our resolution limit of 140-150 nm in activated B cells (Fig. 4a, b). Furthermore, these microclusters moved together towards the center of the contact zone (Supplementary Fig. S4, Supplementary Movie 1). To identify the diffusive states in which CD19 resides, we performed single molecule imaging of CD19 using AF594 labeled anti-CD19 antibody at low labeling concentrations and with the same methods as for single molecule imaging of BCR. By analyzing a collection of CD19 tracks in a control cell during the 10-minute imaging period (Fig. 4c), we found that the diffusivity of CD19 is lower than that of the BCR (compared to Fig. 1d), given the abundance of short tracks (blue) and fewer longer tracks (yellow). pEM analysis of CD19 tracks again resulted in eight distinct states with mean diffusivities preserved over time (Fig. 4d). The pair correlation analysis of CD19 molecules in both control and cNKO cells show higher correlations of states 1 through 4 at shorter distances than the other states, indicative of a clustered configuration (Fig. 4 e, f). As observed for BCRs, trajectories from states 1 and 2 of CD19 show the highest degree of clustering at short distances, while the faster moving states show a more homogeneous distribution.

**Figure 4.**
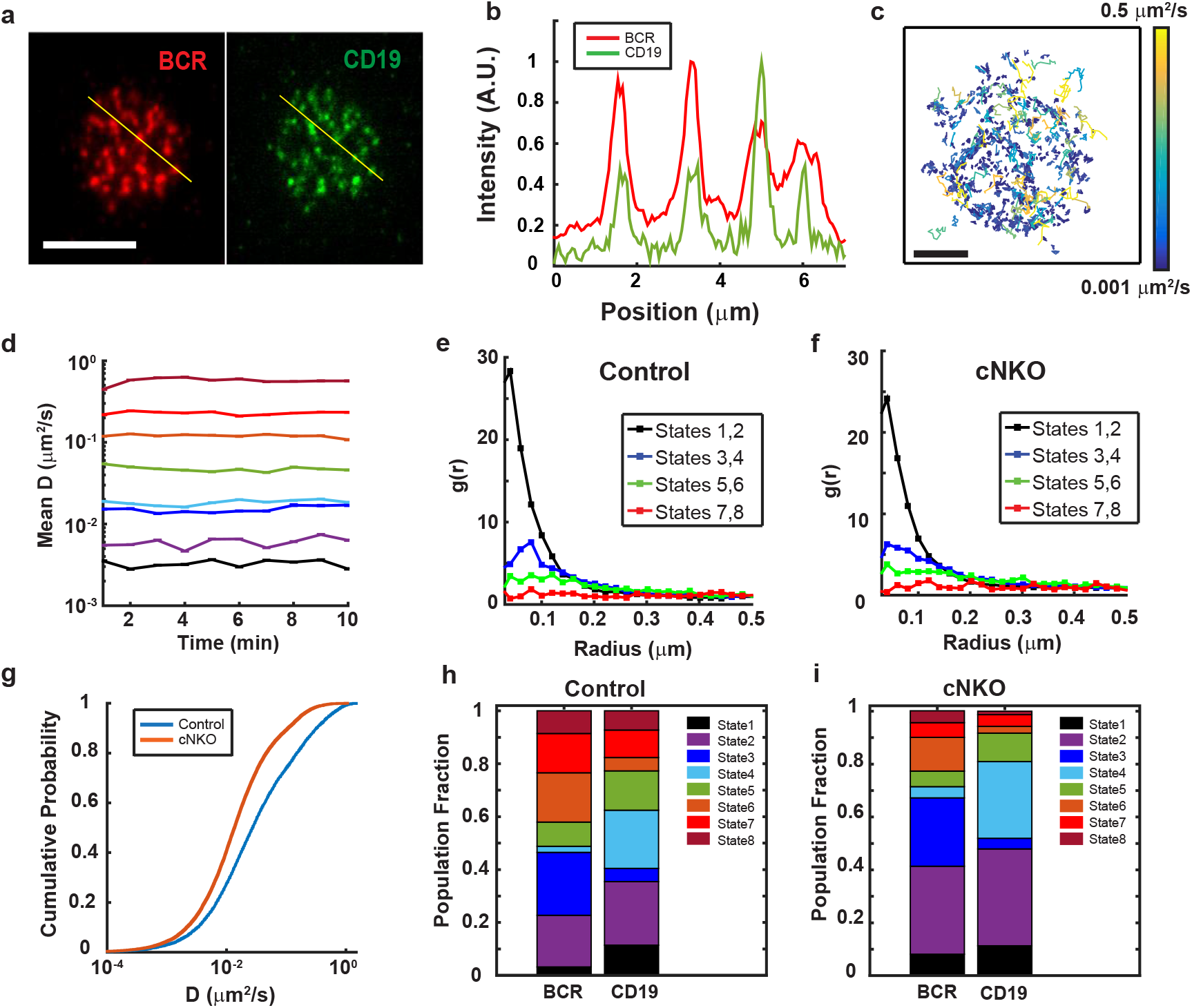
N-WASP expression modulates CD19 diffusivity. **a)** Instant SIM images of activated B cell, showing that AF546 labeled BCR (red) and AF488 labeled CD19 (green) reside in clusters that colocalize to within the ~150 nm resolution limit. Scale bar is 5 μm. **b)** Intensity profiles for BCR (red) and CD19 (green) fluorescence along the yellow lines as drawn in a). **c)** Compilation of CD19 tracks over a 10-minute period in an activated control B cell. Scale bar is 1 μm. **d)** Plot showing the mean diffusivity of each of the eight diffusive states obtained from pEM analysis as a function of time. The colors corresponding to the different states are as shown in Figure 2B. Error bars represent the standard error of the mean. **e, f)** Pair correlation function plot for all states for control and N-WASP-KO cells respectively. **g)** Cumulative Probability distribution of diffusivities showing that mobility of CD19 in cNKO cells is significantly lower than in control cells (control cells N = 10, cNKO cells N = 11, P = 0.0013 Kruskal-Wallis test). **h, i)** Comparison of population fractions of BCR and CD19 in different states for control and cNKO cells respectively.

Interestingly, the cumulative distribution plots of the diffusivities showed that CD19 diffusion in cNKO cells is significantly lower than in control cells (Fig. 4g). The diffusivities of the eight effective states found for CD19 were very similar to those found for BCR, allowing us to compare the population fractions of these states between these two receptors (Fig. 4h). States 1, 2, 4 and 8 are more predominant for CD19 while states 3, 6 and 7 are more populated for BCR in control cells. The population fraction of the lowest mobility states 1 and 2 for CD19 show a significant increase in cNKO cells compared to control cells (Fig. 4i), as observed for BCR. These results suggest that N-WASP knockout out affects the mobility of BCRs and CD19 in a similar way, slowing down their overall mobility and likely maintaining their interactions inside signaling clusters. We thus conclude that BCRs and CD19 in states 1 and 2 are likely to be in a signaling state.

### N-WASP KO has a limited effect on FcγRIIB diffusivity

To determine whether actin-mediated modulation of mobility is specific to BCRs and its stimulatory co-receptor CD19 or reflects a more general change in the diffusive properties of the membrane environment due to changes in the cortical actin network, we tested whether the mobility of FcγRIIB, an inhibitory co-receptor of the BCR, is similarly affected by the lack of N-WASP. The FcγRIIB receptor is a transmembrane receptor that is expressed in B cells and is known to inhibit BCR signaling by the recruitment of phosphatases such as SHIP (SH2-domain containing inositol polyphosphate 5’ phosphatase)^32^ and BCR clustering^33^, when it is colligated with the BCR by antibody-antigen immune complexes. FcγRIIB is known to exhibit relatively high diffusivity in quiescent B cells and its mobility is altered by mutations associated with autoimmune diseases^17^. In the absence of colligation, the mobility of FcγRIIB has the potential to yield important insight into the generic diffusion properties of transmembrane receptors.

We studied the diffusivity of FcγRIIB in activated B cells using the same methods used for the other receptors. Figure 5a shows a compilation of FcγRIIB tracks in a control cell over a 10-minute period. Our analyses showed that in contrast to the significant slowdown of BCR diffusivity in cNKO cells, FcγRIIB diffusivity is minimally affected as shown by the cumulative distribution plot of the diffusivities (Fig. 5b). pEM analysis of the sets of all single molecule trajectories showed 7 states with stable mean diffusivities (Fig. 5c). Pair correlation analysis of tracked FcγRIIB molecules for all states in control and cNKO cells are shown in Figure 5d and e respectively. As with the other receptors studied so far, states 1 and 2 display signs of clustering, states 3 and 5 also display clustering but to a lesser degree, while all other states display a more homogeneous distribution. The population fraction of FcγRIIB in the different diffusive states is similar to BCR, with slight increases in states 1, 2 and 6 (Fig. 5f). As FcγRIIB mobility does not undergo the drastic changes observed in BCR and CD19 mobility in the absence of N-WASP, this result suggests that N-WASP mediated regulation of receptor mobility is highly specific to the BCR.

**Figure 5.**
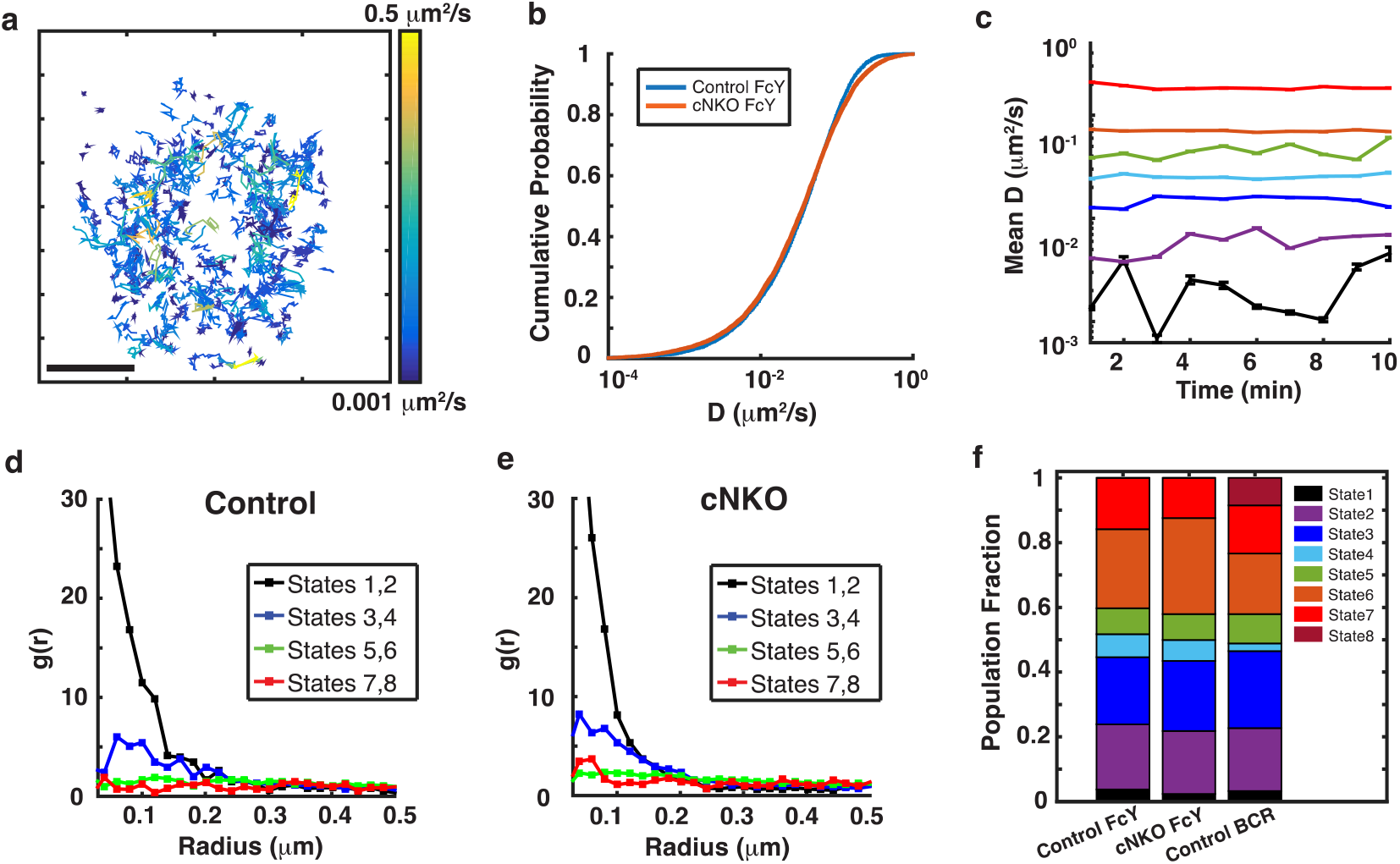
FcγRIIB mobility is mildly affected by the lack of N-WASP. **a)** A set of single molecule tracks of FcγRIIB from an activated B cell over a 10-minute period. Scale bar is 1 μm. b) Cumulative distribution plots for diffusivity of FcγRIIB molecules in control and cNKO cells (control cells N = 12, cNKO cells N = 12). c) pEM analysis of single FcγRIIB molecule trajectories uncovered 7 states. Plot shows the mean diffusivity of each state at every time point. Error bars represent the standard error of the mean. **d, e)** Pair correlation function plot for all states in control and cNKO states respectively. **f)** Comparison of the population fraction of different diffusive states of BCR in control cells and FcγRIIB in control and cNKO cells.

### N-WASP regulates BCR mobility through Arp2/3

N-WASP is known to activate the actin nucleating protein Arp2/3 complex to promote the growth of branched actin networks^34–36^. To investigate whether the effect of cNKO on the mobility of membrane receptors is actin and Arp2/3 mediated, we studied the effect of the small molecule inhibitor of the Arp2/3 complex, CK666 (50 μM), on BCR diffusivity in activated cells. Surface BCRs showed a distinct decrease in mobility in CK666-treated B cells as compared to untreated cells (Fig. 6a), similar to the results from cNKO cells. We next investigated the effect of inhibiting formin, the actin nucleating protein that polymerizes bundled actin, on BCR mobility. B cells treated with the formin inhibitor, SMIFH2 (25 μM) also had BCR with lower mobility as compared to untreated cells (Fig. 6a). The reduction in overall BCR diffusivity caused by formin inhibition was similar to that caused by Arp2/3 inhibition. pEM analysis was performed on the set of BCR tracks from cells treated with these inhibitors. Plots of the population fraction over time of BCRs in B cells treated with CK666 show a trend similar to that observed in cNKO cells, where the low mobility states, states 2 and 3, contribute to over 60% of all trajectories, compared to 40% in control cells (Fig. 6b). The distribution of population fractions for cells treated with SMIFH2 showed a slightly different behavior (Fig. 6c), wherein only state 2 displayed an overall increase (35% of all trajectories) relative to non-treated controls (20% of all trajectories). These results indicate that the effect of cNKO on BCR mobility is likely to be actin driven largely through Arp2/3 mediated branched nucleation, with formin mediated nucleation playing an additional role. We further found that the N-WASP inhibitor wiskostatin (10 μM) also resulted in a decrease in diffusivity but not to the same extent as cNKO (Fig. 6d). Wiskostatin treatment (Fig. 6e) increased the population fraction of BCRs in states 2 and 3, similar to what was observed in cNKO cells but not to the same extent. The inhibition of these actin associated proteins caused a decrease in overall diffusivity of BCRs along with an increase in the fraction of the slow diffusive states as shown in the comparison of population fractions (Fig. 6f). These results collectively implicate actin dynamics in maintaining the heterogeneity of BCR mobility and nanoscale organization.

**Figure 6.**
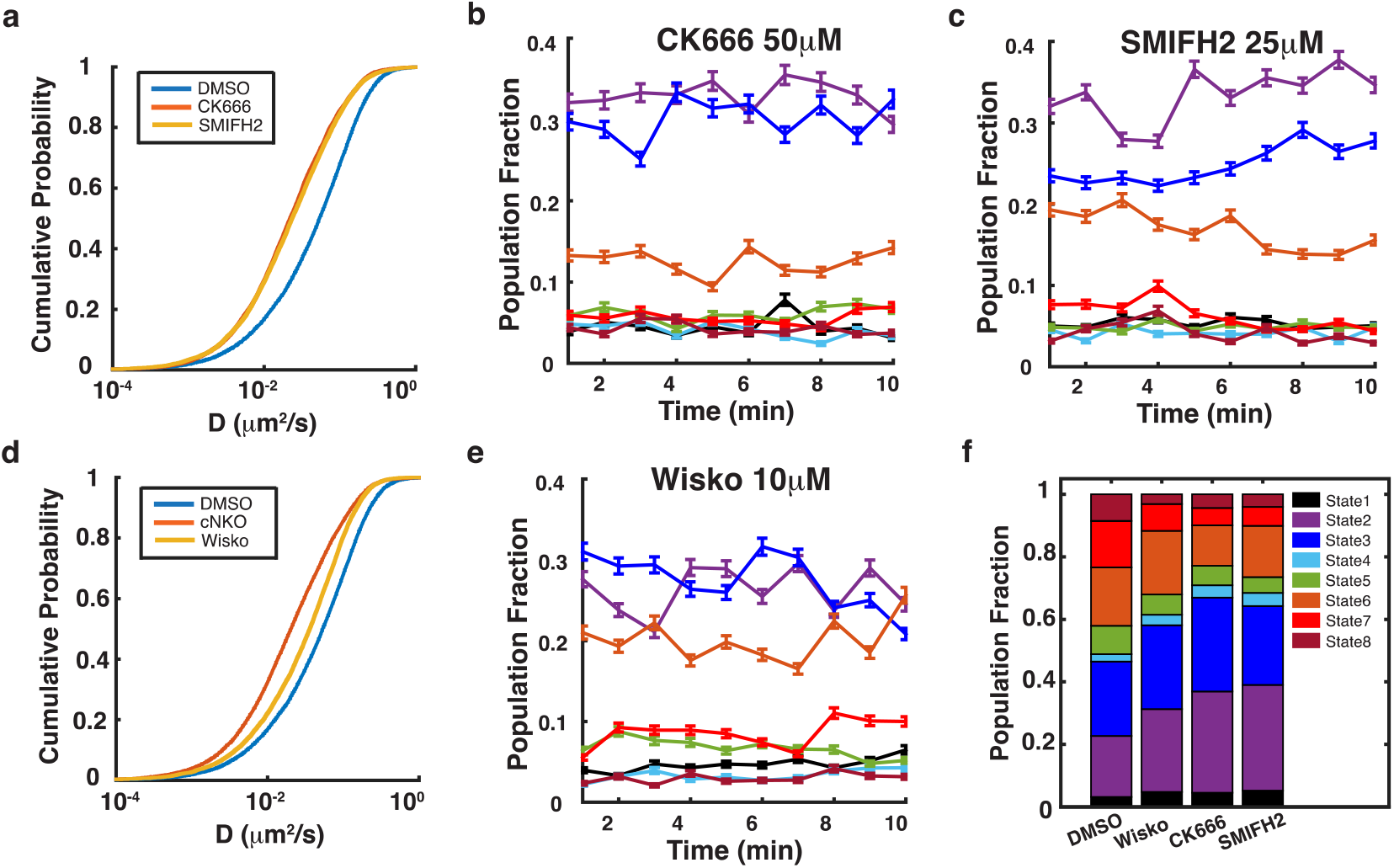
Inhibition of actin nucleation decreases BCR diffusivity. **a)** Plots of BCR diffusivity distributions for cells treated with CK666 or SMIFH2. (P < 0.001, Kruskal-Wallis test for comparison between DMSO and CK666, or DMSO and SMIFH2). **b)** Population fraction over time for cells treated with CK666. **c)** Population fraction over time for cells treated with SMIFH2. **d)** BCR diffusivity distribution for cells treated with wiskostatin (Wisko) compared with cNKO and control. (P < 0.01, Kruskal-Wallis test for comparison between DMSO and Wisko, or control and cNKO) **e)** Population fraction over time for cells treated with wiskostatin. Error bars in B, C and E represent a confidence interval of 95% on the population fraction calculation. **f)** Overall distribution of population fractions for inhibition of N-WASP (Wisko), Arp2/3 complex (CK666) and formin (SMIFH2) (Number of cells: DMSO, N = 14; Wisko, N = 11; CK666, N = 10; SMIFH2, N = 16).

### N-WASP regulates actin dynamics during B cell signaling activation

To obtain insight into how N-WASP driven actin polymerization regulates BCR mobility, we studied actin dynamics in primary B cells from Lifeact-EGFP transgenic mice activated on supported lipid bilayers, and treated with the N-WASP inhibitor, wiskostatin. We used instant structured illumination microscopy (iSIM)^37^ to obtain high spatial resolution images amenable to quantitative analysis for determining actin flows. In primary B cells, the actin network is organized into highly dynamic foci (indicated by blue arrows) and a thin lamellipodial region at the cell periphery (indicated by yellow arrows) in both untreated and wiskostatin treated cells (Fig. 7a, Supplementary Movie 2). We used Spatiotemporal Image Correlation Spectroscopy (STICS)^38^ to estimate the speed and directionality of actin flows (Fig. 7b). We extracted the information from the actin flow velocity vector maps and generated heat maps showing the magnitude of actin speeds and the direction of flow relative to the center of the cell. The magnitude of actin flow speed did not display any systematic spatial dependence in either wiskostatin treated or untreated cells (Fig. 7c). However, wiskostatin treated cells displayed a significant reduction in actin flow speed (Fig. 7d), suggesting that N-WASP is involved in generating a dynamic actin network. To determine the directionality of actin flow vectors, the directional coherence was defined as the cosine of the angle relative to a vector pointing to the centroid of the cell. Thus, flow towards the cell center has value 1 and flow away from the center has value −1 with all other angles spanning intermediate values within this range. The spatial maps of directional coherence values revealed that actin flows were not spatially correlated over large regions of the cell-substrate interface for cells either treated with wiskostatin or untreated (Fig. 7e). We next determined whether the directional coherence measure evolved over time and whether this evolution differed between control and wiskostatin-treated cells. Probability distribution functions of the directional coherence values over the entire contact zone showed that during the early stage of activation (0 - 5 min), actin flows were predominantly directed either inwards or outwards (relative to the cell centroid) with no significant difference between control and wiskostatin-treated cells (Fig. 7f). However, during the late stage of activation (5-10 min), wiskostatin-treated cells displayed greater inward actin flows towards the center of the cell and less outward flows compared to control cells (Fig. 7g).

**Figure 7.**
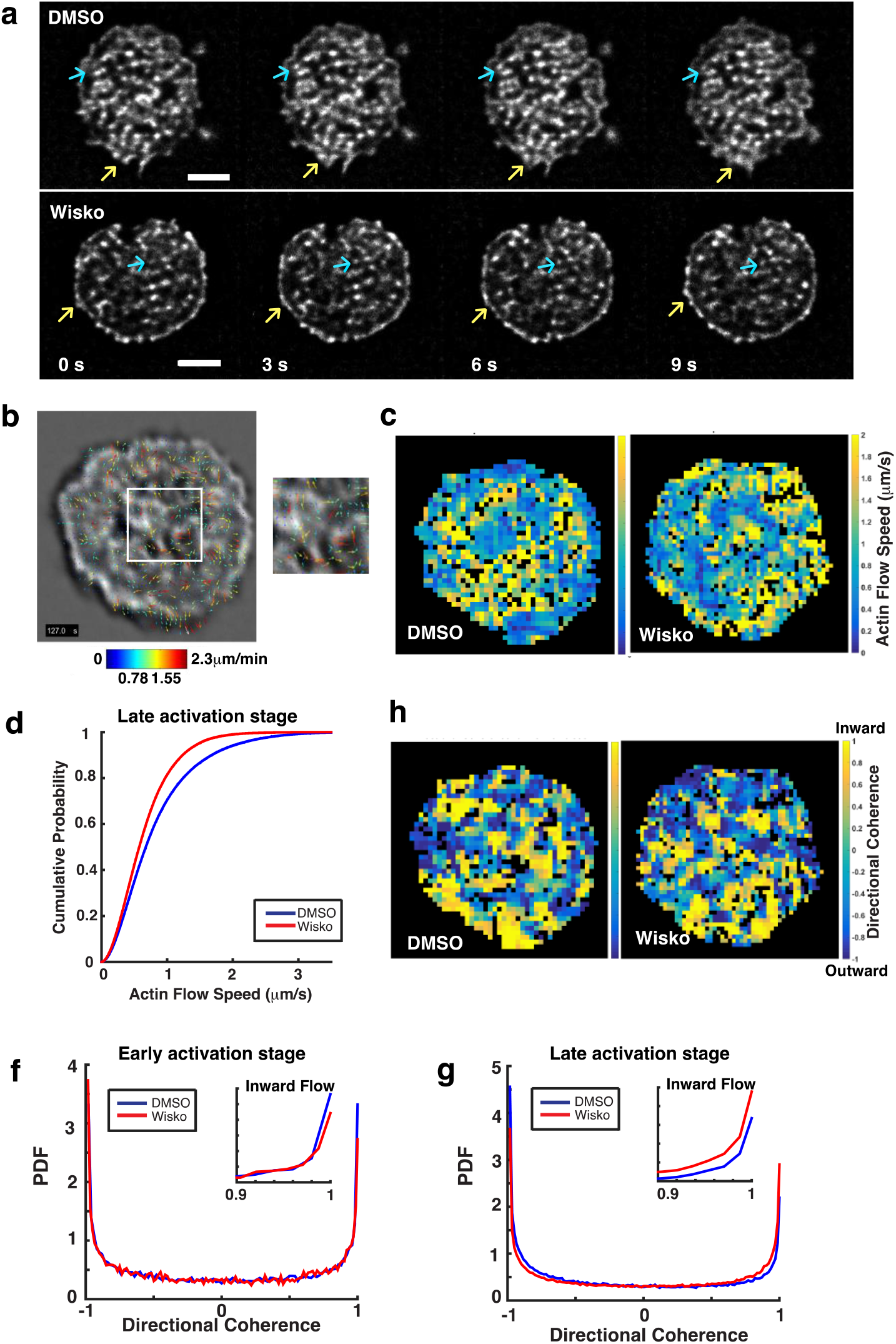
Effect of N-WASP on actin dynamics in activated B cells. **a)** iSIM images of activated Lifeact-EGFP B cells at consecutive time points for two conditions: DMSO carrier-control and N-WASP inhibitor wiskostatin (10 μM concentration). The initial time corresponds to 5 min after spreading initiation. The blue arrows in the images indicate the emergence of actin foci and the yellow arrows point to spreading and contraction of the lamellipodial region of the cells. Scale bars are 2 μm. **b)** STICS (Spatio-Temporal Image Correlation Spectroscopy) vector map showing actin flows represented by velocity vectors indicating flow direction and color coded for flow speed. In the zoomed region, the velocity vectors show the flow direction and flow speed. The vector map is overlaid on top of a grayscale image of Lifeact-EGFP. **c)** Pseudocolor map of actin flow speeds corresponding to 2 minutes after cell spreading for representative DMSO control and wiskostatin treated cells. **d)** Cumulative distribution of actin flow speeds for DMSO control cells (blue, N = 11 cells) and cells treated with wiskostatin (red, N = 12 cells). (P = 0.00074, Kruskal-Wallis test). e) Directional coherence maps indicating the flow directions, which ranged from inward (1) to outward (-1). **f, g)** Probability distribution function plots showing directional coherence values of actin flow in cells during the early stage of activation **f)** or cells in the late stage of activation **g)**, with subplots highlighting the flow fraction defined as inward flow (see methods). During early stage, fraction of inward flow is 0.143 for DMSO and 0.1425 for Wisko treated cells (P = 0.5188 - not significant); during late stage the fraction of inward flow is 0.113 for DMSO and 0.1518 for Wisko treated cells (P < 0.001).

## Discussion

Here, we used single molecule imaging to study how actin dynamics regulates BCR diffusion and thereby signaling initiation by examining the role of the actin regulatory protein N-WASP in regulating the diffusivity of surface BCRs and its stimulatory and inhibitory co-receptors. Using perturbation Expectation Maximization, a method that analyzes receptor trajectories at the single molecule level, we found that surface BCRs exist in distinct diffusive states. The population of the fastest state, which likely corresponds to monomeric BCR, is reduced over the first 4 minutes of stimulation. Using pair correlation analysis to quantify the degree of clustering exhibited by BCRs and relate them to their local mobility, we found that states with low diffusivities display a greater degree of clustering. Our studies further showed that BCRs in lower mobility states are significantly more predominant in cNKO cells. Given previous studies showing that N-WASP knockout is associated with enhanced signaling^18^, our observations suggest that low mobility BCR trajectories are associated with signaling states. Our findings reveal a role for actin regulatory proteins in modulating receptor dynamics, highlighting the importance of the dynamic actin network in regulating receptor mobility and signaling.

Using both pair correlation and pEM analysis, we found that the low diffusivity of BCRs on the surface of cNKO B cells is accompanied by a lower diffusivity of its stimulatory co-receptor CD19, again strengthening our hypothesis that signaling states of BCRs correlate with those that display decreased mobility. However, lack of N-WASP did not affect the diffusivity of FcγRIIB, an inhibitory co-receptor of the BCR, suggesting that its effect on BCR and CD19 mobility is highly specific to signaling activation. Our findings are broadly consistent with prior studies. Stone et al. showed that the spatial positions of the BCR and Lyn, a signaling kinase, become correlated after antigen stimulation, and this correlation is accompanied by a reduction in diffusivity of both molecules^39^. Using a photoactivatable antigen, Wang et al.^40^ measured the diffusivity of BCR on the same set of cells before and after activation and found a decrease in BCR diffusivity following antigenic stimulation. During activation, conformational changes of ITAM-containing receptors, changes in local lipid environment and interactions with other proteins may alter the mobility of membrane receptors. For instance, stimulation of mast cells through FcεRI receptor crosslinking, induces the clustering of FcεRI and a concomitant reduction of its diffusivity that depends on the average number of receptors in the cluster^41^. The clustering of FcεRI receptor is accompanied by the redistribution of the signaling proteins Lyn kinase and Syk kinase into clusters^42^.

Inhibition of the Arp2/3 complex, which nucleates branched actin networks downstream of N-WASP, also resulted in an overall reduction of BCR diffusivity. This is consistent with previous work, which has shown that inhibition, or knockout of N-WASP induces higher and more prolonged signaling. Moreover, the reduction in BCR diffusivity by Arp2/3 inhibition was similar to what was observed in cNKO B cells, suggesting that the primary effect of N-WASP is mediated through activation of Arp2/3-mediated actin nucleation. Another important class of actin nucleators are formins, which assemble bundled actin structures. Interestingly, inhibition of formin also resulted in reduced BCR mobility, but with somewhat different effects on the diffusive states as compared to the effects of Arp2/3 inhibition. Previous studies both in fission yeast and mammalian cells^43,44^ have suggested that the two nucleators compete for actin monomers, and that the disruption of either leads to increased activity of the other and the formation of distinct actin networks. Our findings show that disruption of both Arp2/3 and formin mediated dynamic actin networks affects receptor mobility. In summary, our studies suggest that optimal BCR signaling requires homeostatic balance between actin networks generated by multiple nucleating proteins.

A well-accepted model of the regulation of BCR diffusion by the actin cytoskeleton in resting B cells posits that the actin network imposes diffusional barriers on BCR and other receptors and signaling proteins (Fig. 8a). Activation leads to the dissolution of these barriers, thereby enhancing BCR diffusion and leading to activation. Early BCR signaling also leads to increased actin dynamics (Fig. 8b). Our imaging studies have revealed that B cell actin remodeling is not characterized by spatially coherent directional flows. Rather, actin dynamics is highly complex, with sharp changes in speed and directionality, both spatially and temporally. This dynamics may also be associated with the formation of non-equilibrium actin structures such as asters and foci^45,46^. We propose that these rapidly changing actin flows may serve to ‘stir’ the cytoplasm adjacent to the membrane, thus changing the reaction environment of receptors and signaling molecules, and modulating the reaction rates in the juxta-membrane regions of the cytoplasm^47^. Combined with the release of BCRs from diffusion traps, this active stirring may drive receptors into clusters and facilitate receptor interactions with activating kinases^19^. At later stages, further increases in actin dynamics and increased outward actin flows could decrease reaction rates between BCRs and activating kinases or increase reaction rates between BCRs and inhibitory co-receptors, making signaling states more unstable, facilitating down-regulation of signaling (Fig. 8c top). When N-WASP is inhibited or knocked out (Fig. 8c bottom), B cells exhibit reduced actin dynamics, which may decrease actin-mediated mixing in the membrane, likely enabling BCR to enter and remain in signaling states (clusters) leading to enhanced signaling and preponderance of low mobility states (or conversely increased interactions with phosphatases leading to signaling inhibition).

**Figure 8.**
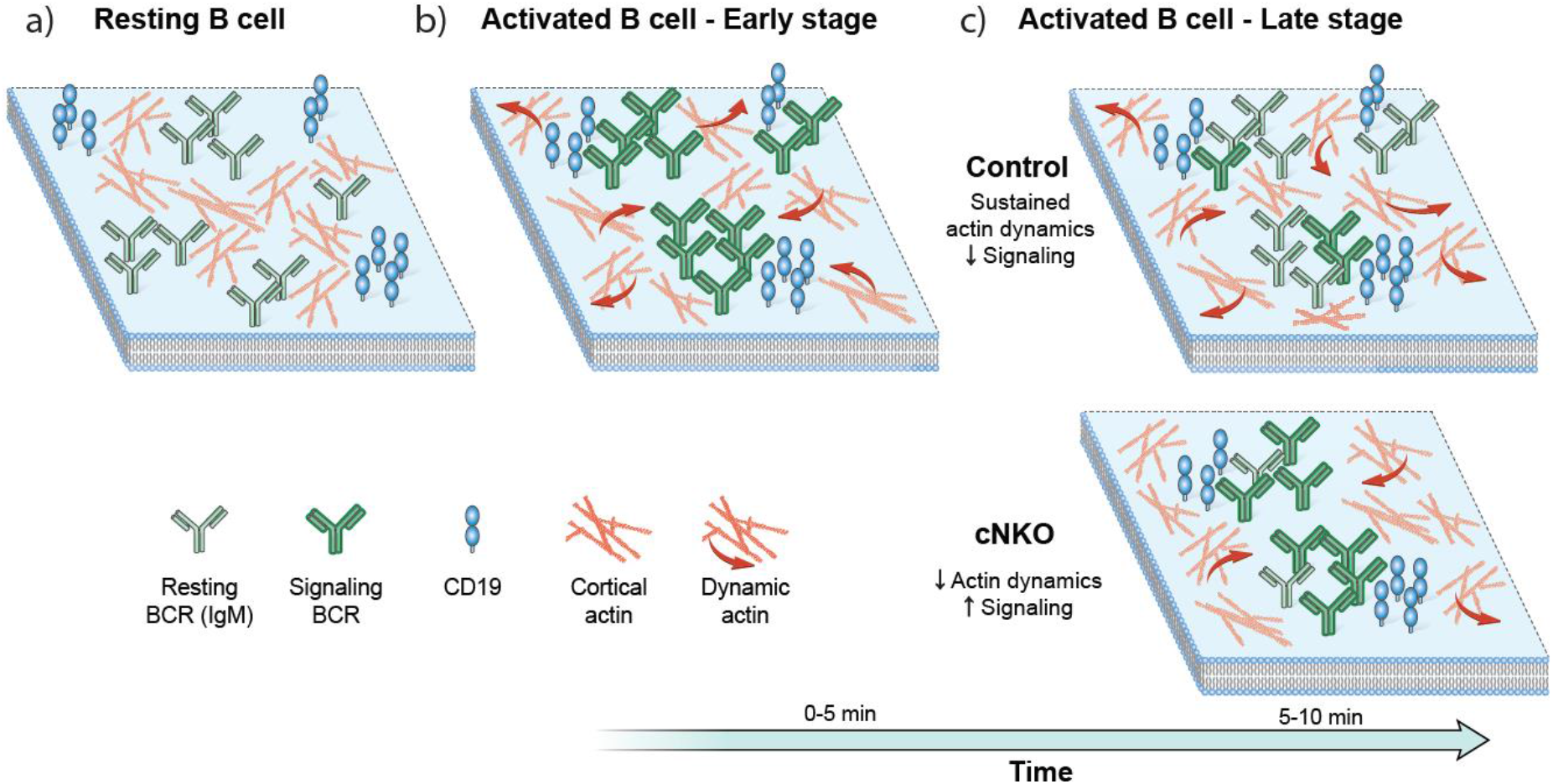
The actin cytoskeleton regulates B cell receptor mobility and signaling in different stages. Representative cartoon showing receptor distributions on a section of the B cell membrane: **a)** Resting B cell membrane: Actin networks restrict receptor lateral movement and interactions. **b)** B cell membrane at the early signaling activation stage. Actin remodeling enhances receptor mobility allowing for interactions between receptors, specifically BCR and CD19, enhancing signaling. Actin flows towards the center and edge of the immune synapse in similar proportions. **c)** B cell membrane at later activation stages. Top: Actin flows stir the cytoplasm at the membrane vicinity, increasing the mixing of receptors in the membrane and thereby allowing signal inhibitory molecules to downregulate BCR signaling. Bottom: N-WASP knockout reduces actin dynamics and changes the balance of actin flow directionality at later stages (5-10 min) of activation, leading to enhanced signaling.

Based on our observations, we suggest that actin dynamics in the cell may be used to fine-tune the levels of signaling activation. Modulation of the composition and structure of actin networks, by changing the expression levels or spatial distribution of actin regulatory proteins, may provide the cell with a powerful way to regulate signaling over rapid timescales. These properties are likely to be a general feature of cells in the immune system whose function depends on rapid response to external stimuli, and illustrate general principles of ITAM receptor signaling.

## Methods

### Mice and cell preparation

B-cell–specific N-WASP knockout (CD19^Cre/+^ N-WASP^Flox/Flox^, cNKO) mice and control mice (CD19^+/+^ N-WASP^Flox/Flox^) were bred as previously described^18^. Transgenic Lifeact-EGFP mice are as described before^48^. Mice selected for experiments were between 2 to 4 months old with no gender preference. Naïve primary B cells were isolated from mouse spleens using a negative selection procedure as described before^23^. After extraction, cells were kept at 4 °C and cell aliquots were pre-warmed at 37 °C for 5 minutes before being added to the bilayer. All experiments involving animals have been approved by the University of Maryland Institution Animal Care and Usage Committee.

### Fluorescent antibodies and inhibitors

For inhibition of formin and Arp2/3, cells were incubated with inhibitors for 5 minutes at 37 °C before being added to the imaging chamber, which had the inhibitor at the same concentration used for incubation. SMIFH2 (Sigma-Aldrich) was used at a 25 μM concentration. Arp2/3 complex inhibitor I, CK-666 (Calbiochem) was used at 50 μM. For N-WASP inhibition, wiskostatin B (EMD Bioscience) was used at 10 μM to incubate the cells for 1 hour at 37 °C. Mono-biotinylated fragment of antibody (mbFab’-anti-Ig) was generated from the F(ab’)_2_ fragment (Jackson Immuno Research, West Grove PA) using a published protocol^49^. FcγRIIB (CD32) antibody (BD Biosciences) was conjugated with Alexa Fluor 546 using Molecular Probes Protein labeling kits (Invitrogen) following manufacturer protocols. For labeling of CD19 we used the Alexa Fluor 594 anti mouse CD19 antibody (BioLegend).

### Sample preparation for single particle tracking

Glass slides were kept in nanostrip overnight and then rinsed with dd-H2O and dried with filtered air. Supported lipid bilayers were prepared by incubating slides with 10 μM DOPC/DOPE-cap-biotin liposome solution for 10 minutes at room temperature. The slides were rinsed with filtered PBS (1X) and then incubated for 10 minutes with 1 μg/ml solution of streptavidin. Slides were rinsed again with PBS and then incubated with unlabeled mono-biotinylated fragment of antibody (mbFab) solution at 18 μg/ml. PBS was replaced with L-15 (CO_2_ independent media with 2% FBS) before imaging. 0.75 μl of 0.05 mg/ml AF546 labeled mbFab was added to a 250 μl volume of media in the imaging chamber.

### Microscopy

For single molecule imaging of BCR we used an inverted microscope (Nikon TE2000 PFS) equipped with a 1.49 NA 100X lens for TIRF imaging and an EMCCD camera (iXon 897, Andor). In order to image single molecules on the cell membrane for extended periods of time we add a low concentration of the fluorescent antibody in solution as shown in Figure 1. Cells are imaged from the moment they reach the bilayer and time-lapse movies (1000 frames acquired at 33 Hz) representative of each minute are taken. Figure 1B shows a representative frame where the cell outline is obtained from an IRM image taken after the single molecule movie. The molecules detected at each frame are localized with high precision (~20 nm) and linked frame by frame to create tracks^27^ using a MATLAB routine. Taking into account motion blur, pEM estimates localization precision to range between 20 nm (slowest states) to 80 nm (fastest state). Imaging of Lifeact-EGFP expressing murine primary cells spreading on supported lipid bilayers was performed using instant Structured Illumination Microscopy (iSIM)^37^, with a 60X 1.42 N.A. lens (Olympus), a 488 nm laser for excitation with 200 ms exposure times and a PCO Edge camera. Images obtained are post-processed with background subtraction and deconvolution. The final lateral resolution for deconvolved images is between 140-150 nm. Spreading cells were imaged at 2 second intervals and spread cells were imaged at 5 frames per second. The Richardson-Lucy algorithm is used for deconvolution, and run for 10 iterations. The PSF used was simulated by a Gaussian function but based on parameters obtained from measurement, i.e. the FWHM of the PSF used is the same as the FWHM measured.

### Data analysis

Perturbation-Expectation maximization version 2 (pEM v2) was used to classify single molecule tracks derived from different receptors^26^. To perform pEM analysis all tracks must have the same length. Given the 33 Hz imaging rate, the optimal track length was found to be 15 frames long due to the tradeoff between accurately identifying diffusivities and minimizing the number of state transitions that the particle may undergo over a single trajectory^50^. All single molecule trajectories obtained were split into 15 frame segments and the classification analysis was performed on the set of all these track segments. Trajectories larger than 105 frames or shorter than 15 frames were discarded. The tracking routine interrupts the creation of a trajectory whenever two particles cross paths. To avoid an over counting of slow-moving molecules (which have lower probability of crossing paths with other molecules) we discarded trajectories longer than 105 frames. The data was then separated according to the receptor type and PEM v2 was run for all data sets using 20 re-initializations, 150 perturbations, 14 covariance parameters and allowing the system to explore up to 15 states. This set of parameter values was chosen to ensure convergence to the global maximum. For all conditions, the average track length was 40 frames and typically 100 tracks were obtained per cell per time point. The maximum posterior probability value was used to assign a track uniquely to a particular state as shown in Supplementary Figure S5A for BCR in control cells. For BCR, 186959 tracks corresponding to all inhibitor treatments, DMSO control and cNKO cells were analyzed together. For CD19, 35062 tracks corresponding to control and cNKO were pooled together and analyzed, and for FcγRIIB receptor, 24969 tracks were analyzed. For all the receptors and conditions 8 diffusive states were identified. The states were compared across different receptors based on a comparison of their diffusivity distributions (Supplementary Figure S5B).

STICS analysis of actin flows was implemented on iSIM images taken at 2 second intervals. Sub-regions of 8 x 8 pixels were selected with a shift of 2 pixels between sub-regions. Immobile filtering was set to 20 frames and the time of interest (TOI) was chosen as 5 frames with a shift of 3 frames between TOIs. Velocity flow vectors that exceeded the sub-region threshold were discarded, giving place to the ‘black pixels’ observed in the decomposed maps of speeds and directions. To determine the directionality of the flow, the centroid of the cell was calculated and a vector from each of the sub-regions pointing towards the centroid was obtained. The directional coherence was then determined as the cosine of the angle between velocity vector and the vector pointing to the centroid. Directionality plots were generated using the MATLAB function histcounts and using probability density function (PDF) as normalization type. In order to compare directionality between DMSO and wiskostatin treated cells the fraction of inward flow (values larger than 0.9) and the fraction of outward flow (values less than -0.9) was determined. The comparison of fraction of flow in either direction between the two conditions was tested using the Z test where the null hypothesis is that both fractions are equal.

For receptor diffusivity studies 12 control and 9 cNKO mice were used. For actin dynamics studies 3 Lifeact-EGFP mice were used.

### Statistical analysis

The Kruskal-Wallis test was used to assess the difference between the diffusivity distributions corresponding to different conditions. We used this test for most comparisons because is a non-parametric method for testing whether two data samples originate from the same distribution. The test was performed over smaller data subsets selected randomly and implemented using the kruskalwallis function in MATLAB. The pair wise Z test was used to determine the difference in proportions of diffusive states across different conditions.

#### Code availability

The code for pEM analysis is freely available at the following link: https://github.com/p-koo/pEMv2

## Acknowledgements

A.U. acknowledges support from the grant NSF PHY 1607645. A.U. and W.S. acknowledge support from the grant NIH R21 AI122205. H.S. acknowledges support from the Intramural Research Programs of the National Institute of Biomedical Imaging and Bioengineering. S.M acknowledges support by NIGMS AWDA 10958 and the NSF Physics of Living Systems Student Research Network under PHYS 1522467. I. R-S would like to acknowledge support from the Fulbright-Colciencias scholarship. We would like to thank Dr. Peter Tuo Li and Dr. Tom Blanpied (UMD, School of Medicine) for guidance on the design of the single molecule tracking experiments. We thank Harsh Vishwasrao and Jiji Chen of the NIH Advanced Imaging and Microscopy resource for help with iSIM and Kaustubh Wagh (UMD) for useful discussions on actin dynamics analysis.

## Author contributions

I.R.-S. and A.U. designed the experiments with input from W.S. I.R.-S. and B.W. performed the experiments and analyzed the data. P.K. and S.M. provided guidance on PEM analysis. Z.S. and W.S. provided guidance on spleen and bilayer preparation. H.S. provided guidance on imaging and analysis. I.R.-S. and A.U. wrote the original draft of the manuscript, with input and editing from H.S., S.M. and W.S. A.U. supervised the project.

## Data Availability

All the data is available upon reasonable request.

## References

1 Fleire, S. J. et al. B cell ligand discrimination through a spreading and contraction response. Science 312, 738–741.

2 Kuokkanen, E., Sustar, V. & Mattila, P. K. Molecular control of B cell activation and immunological synapse formation. Traffic 16, 311–326.

3 Cambier, J. C., Pleiman, C. M. & Clark, M. R. Signal transduction by the B cell antigen receptor and its coreceptors. Annu Rev Immunol 12, 457–486.

4 Batista, F. D., Iber, D. & Neuberger, M. S. B cells acquire antigen from target cells after synapse formation. Nature 411, 489–494.

5 Rolli, V. et al. Amplification of B cell antigen receptor signaling by a Syk/ITAM positive feedback loop. Molecular cell 10, 1057–1069.

6 Sohn, H. W., Tolar, P., Jin, T. & Pierce, S. K. Fluorescence resonance energy transfer in living cells reveals dynamic membrane changes in the initiation of B cell signaling. Proc Natl Acad Sci U S A 103, 8143–8148.

7 Sohn, H. W., Tolar, P. & Pierce, S. K. Membrane heterogeneities in the formation of B cell receptor-Lyn kinase microclusters and the immune synapse. J Cell Biol 182, 367–379.

8 Tolar, P., Sohn, H. W., Liu, W. & Pierce, S. K. The molecular assembly and organization of signaling active B-cell receptor oligomers. Immunological reviews 232, 34–41.

9 Tolar, P., Hanna, J., Krueger, P. D. & Pierce, S. K. The constant region of the membrane immunoglobulin mediates B cell-receptor clustering and signaling in response to membrane antigens. Immunity 30, 44–55.

10 Charrier, C., Ehrensperger, M. V., Dahan, M., Levi, S. & Triller, A. Cytoskeleton regulation of glycine receptor number at synapses and diffusion in the plasma membrane. The Journal of neuroscience: the official journal of the Society for Neuroscience 26, 8502–8511.

11 Lenne, P. F. et al. Dynamic molecular confinement in the plasma membrane by microdomains and the cytoskeleton meshwork. EMBO J25, 3245–3256.

12 Sheetz, M. P., Schindler, M. & Koppel, D. E. Lateral mobility of integral membrane proteins is increased in spherocytic erythrocytes. Nature 285, 510–511.

13 Treanor, B. et al. The membrane skeleton controls diffusion dynamics and signaling through the B cell receptor. Immunity 32, 187–199.

14 Treanor, B., Depoil, D., Bruckbauer, A. & Batista, F. D. Dynamic cortical actin remodeling by ERM proteins controls BCR microcluster organization and integrity. J Exp Med 208, 1055–1068.

15 Freeman, S. A. et al. Toll-like receptor ligands sensitize B-cell receptor signalling by reducing actin-dependent spatial confinement of the receptor. Nature communications 6, 61–68.

16 Gasparrini, F. et al. Nanoscale organization and dynamics of the siglec CD22 cooperate with the cytoskeleton in restraining BCR signalling. EMBO J 35, 258–280.

17 Xu, L. et al. Impairment on the lateral mobility induced by structural changes underlies the functional deficiency of the lupus-associated polymorphism FcgammaRIIB-T232. J Exp Med 213, 2707–2727.

18 Liu, C. et al. N-wasp is essential for the negative regulation of B cell receptor signaling. PLoS Biol 11, e1001704.

19 Tolar, P. Cytoskeletal control of B cell responses to antigens. Nat Rev Immunol 17, 621–634.

20 Ketchum, C., Miller, H., Song, W. & Upadhyaya, A. Ligand mobility regulates B cell receptor clustering and signaling activation. Biophys J 106, 26–36.

21 Brownlie, R. J. et al. Distinct cell-specific control of autoimmunity and infection by FcgammaRIIb. J Exp Med 205, 883–895.

22 Fearon, D. T. & Carroll, M. C. Regulation of B lymphocyte responses to foreign and self-antigens by the CD19/CD21 complex. Annu Rev Immunol 18, 393–422.

23 Rey, I., Garcia, D. A., Wheatley, B. A., Song, W. & Upadhyaya, A. Biophysical Techniques to Study B Cell Activation: Single-Molecule Imaging and Force Measurements. Methods in molecular biology 1707, 51–68.

24 Frost, N. A., Shroff, H., Kong, H., Betzig, E. & Blanpied, T. A. Single-molecule discrimination of discrete perisynaptic and distributed sites of actin filament assembly within dendritic spines. Neuron 67, 86–99.

25 Vestergaard, C. L., Blainey, P. C. & Flyvbjerg, H. Optimal estimation of diffusion coefficients from single-particle trajectories. Physical review 89, 022726.

26 Koo, P. K. & Mochrie, S. G. J. Applying Perturbation Expectation-Maximization to Protein Trajectories of Rho GTPases. Methods in molecular biology 1821, 57–70.

27 Koo, P. K., Weitzman, M., Sabanaygam, C. R., van Golen, K. L. & Mochrie, S. G. Extracting Diffusive States of Rho GTPase in Live Cells: Towards In Vivo Biochemistry. PLoS Comput Biol 11, e1004297.

28 Sengupta, P. et al. Probing protein heterogeneity in the plasma membrane using PALM and pair correlation analysis. Nature methods 8, 969–975.

29 Depoil, D. et al. CD19 is essential for B cell activation by promoting B cell receptor-antigen microcluster formation in response to membrane-bound ligand. Nat Immunol 9, 63–72.

30 Maity, P. C. et al. B cell antigen receptors of the IgM and IgD classes are clustered in different protein islands that are altered during B cell activation. Sci Signal 8, ra93.

31 Mattila, P. K. et al. The actin and tetraspanin networks organize receptor nanoclusters to regulate B cell receptor-mediated signaling. Immunity 38, 461–474.

32 Ono, M., Bolland, S., Tempst, P. & Ravetch, J. V. Role of the inositol phosphatase SHIP in negative regulation of the immune system by the receptor Fc(gamma)RIIB. Nature 383, 263–266.

33 Xu, L. et al. Through an ITIM-independent mechanism the FcgammaRIIB blocks B cell activation by disrupting the colocalized microclustering of the B cell receptor and CD19. J Immunol 192, 5179–5191.

34 Machesky, L. M. & Insall, R. H. Scar1 and the related Wiskott-Aldrich syndrome protein, WASP, regulate the actin cytoskeleton through the Arp2/3 complex. Curr Biol 8, 1347–1356.

35 Miki, H., Miura, K. & Takenawa, T. N-WASP, a novel actin-depolymerizing protein, regulates the cortical cytoskeletal rearrangement in a PIP2-dependent manner downstream of tyrosine kinases. EMBO J 15, 5326–5335.

36 Symons, M. et al. Wiskott-Aldrich syndrome protein, a novel effector for the GTPase CDC42Hs, is implicated in actin polymerization. Cell 84, 723–734.

37 York, A. G. et al. Instant super-resolution imaging in live cells and embryos via analog image processing. Nature methods 10, 1122–1126.

38 Hebert, B., Costantino, S. & Wiseman, P. W. Spatiotemporal image correlation spectroscopy (STICS) theory, verification, and application to protein velocity mapping in living CHO cells. Biophys J 88, 3601–3614.

39 Stone, M. B., Shelby, S. A., Núñez, M. F., Wisser, K. & Veatch, S. L. Protein sorting by lipid phase-like domains supports emergent signaling function in B lymphocyte plasma membranes. eLife 6, e19891.

40 Wang, J. et al. Utilization of a photoactivatable antigen system to examine B-cell probing termination and the B-cell receptor sorting mechanisms during B-cell activation. Proc Natl Acad Sci US A 113, E558–567.

41 Veatch, S. L., Chiang, E. N., Sengupta, P., Holowka, D. A. & Baird, B. A. Quantitative nanoscale analysis of IgE-FcepsilonRI clustering and coupling to early signaling proteins. The journal of physical chemistry 116, 6923–6935.

42 Shelby, S. A., Veatch, S. L., Holowka, D. A. & Baird, B. A. Functional nanoscale coupling of Lyn kinase with IgE-FcepsilonRI is restricted by the actin cytoskeleton in early antigen-stimulated signaling. Molecular biology of the cell 27, 3645–3658.

43 Burke, T. A. et al. Homeostatic actin cytoskeleton networks are regulated by assembly factor competition for monomers. Curr Biol 24, 579–585.

44 Lomakin, A. J. et al. Competition for actin between two distinct F-actin networks defines a bistable switch for cell polarization. Nat Cell Biol 17, 1435–1445.

45 Fritzsche, M. et al. Cytoskeletal actin dynamics shape a ramifying actin network underpinning immunological synapse formation. Science advances 3, e1603032.

46 Kumari, S. et al. Actin foci facilitate activation of the phospholipase C-gamma in primary T lymphocytes via the WASP pathway. eLife 4.

47 Chaudhuri, A., Bhattacharya, B., Gowrishankar, K., Mayor, S. & Rao, M. Spatiotemporal regulation of chemical reactions by active cytoskeletal remodeling. Proc Natl Acad Sci U S A 108, 14825–14830.

48 Riedl, J. et al. Lifeact mice for studying F-actin dynamics. Nature methods 7, 168–169.

49 Peluso, P. et al. Optimizing antibody immobilization strategies for the construction of protein microarrays. Analytical biochemistry 312, 113–124.

50 Koo, P. K. & Mochrie, S. G. Systems-level approach to uncovering diffusive states and their transitions from single-particle trajectories. Physical review. E 94, 052412.

