## Supplementary Fig. for "N-WASP regulates the mobility of the B cell receptor and co-receptors during signaling activation"

**Supplementary Movie 1.** Time lapse TIRF images showing the coalescence of clusters and their inward motion in a B cell 2 minutes after spreading initiation on a bilayer. The panels show AF546 labeled mbFab (labeling BCR) in red, AF488 labeled anti-CD19 in green, and the composite of both channels showing the movement and colocalization of clusters.

**Supplementary Movie 2.** Time lapse images showing actin dynamics in B cells from Lifeact-EGFP mice for DMSO control and Wiskostatin treated cells.

**Supplementary Table 1.** Percent change and significance of the difference (from z test) for the population fraction of each state in cNKO cells compared with control cells. A positive value indicates an increase in the population fraction of a state in cNKO condition.

| | **State 1** | **State 2** | **State 3** | **State 4** | **State 5** | **State 6** | **State 7** | **State 8** |
|---|---|---|---|---|---|---|---|---|
| **Percent change** | 153.14 | 70.61 | 8.46 | 75.61 | -35.72 | -31.60 | -62.69 | -46.99 |
| **P** | < 0.001 | < 0.001 | 0.01500 | 0.0324 | < 0.001 | < 0.001 | < 0.001 | < 0.001 |

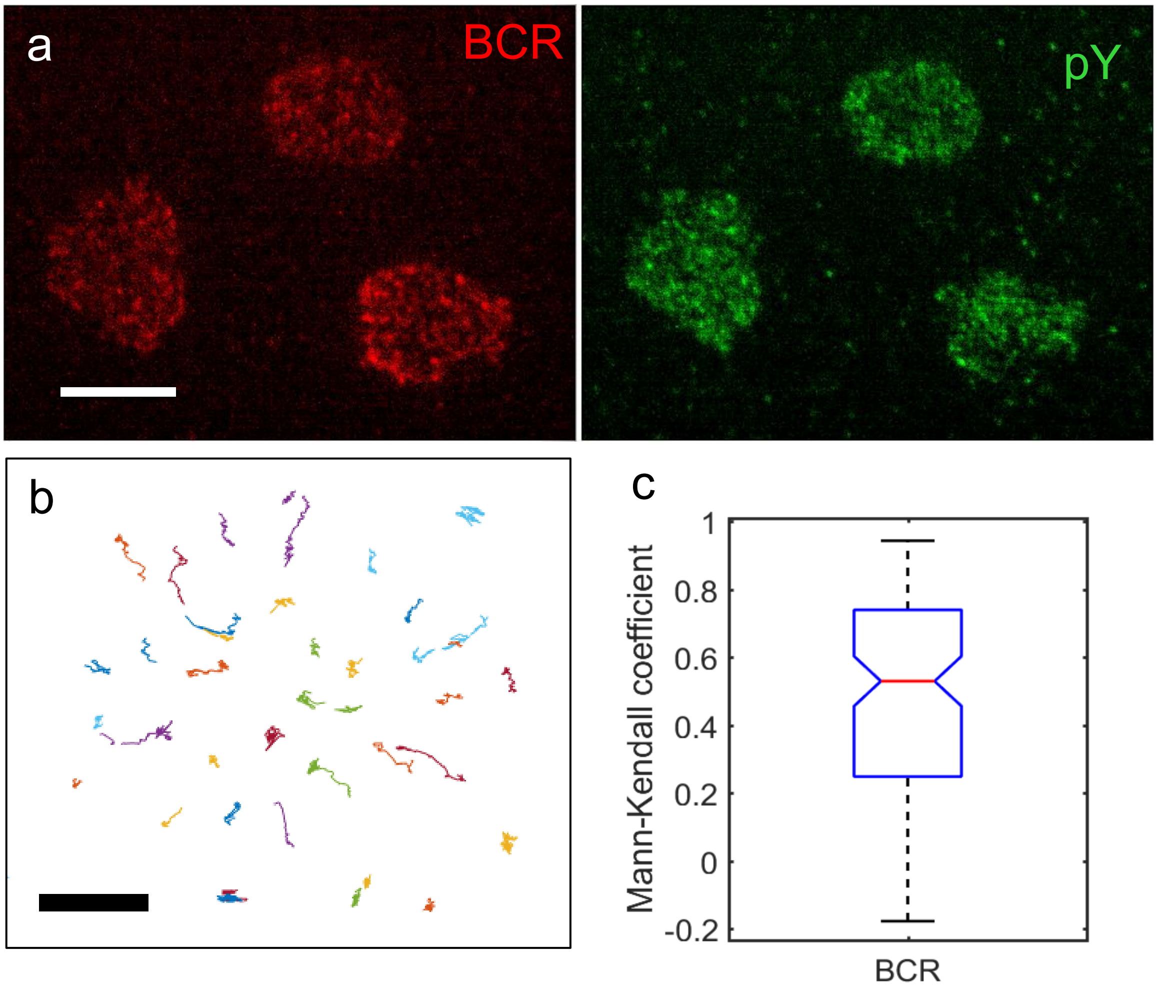


**Supplementary Figure S1. a)** iSIM images of fixed cells (fixed after 3 min of spreading initiation) activated under the same conditions as the single molecule experiments. The red color corresponds to AF546 BCR labeled microclusters while the green is AF488 labeled phosphorylated tyrosine, which also accumulated in clusters. Scale bar is 5 µm. b) BCR microlusters were tracked as they moved towards the center of the cell during the first 200 seconds after the cell contacted the bilayer. Scale bar is 1 µm. c) The intensity within clusters over time was quantified (N = 10 cells) and the Mann-Kendall coefficient was calculated to show the overall increase in intensity over time (MK > 0).

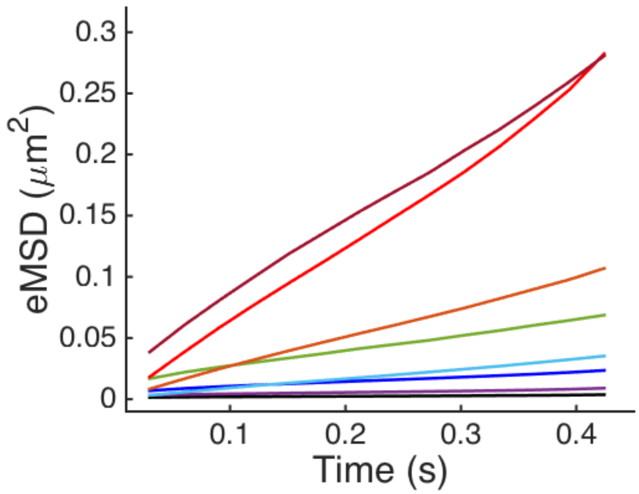


**Supplementary Figure S2.** Ensemble mean square displacement (eMSD) plots for the 8 states identified for BCR in cNKO cells.

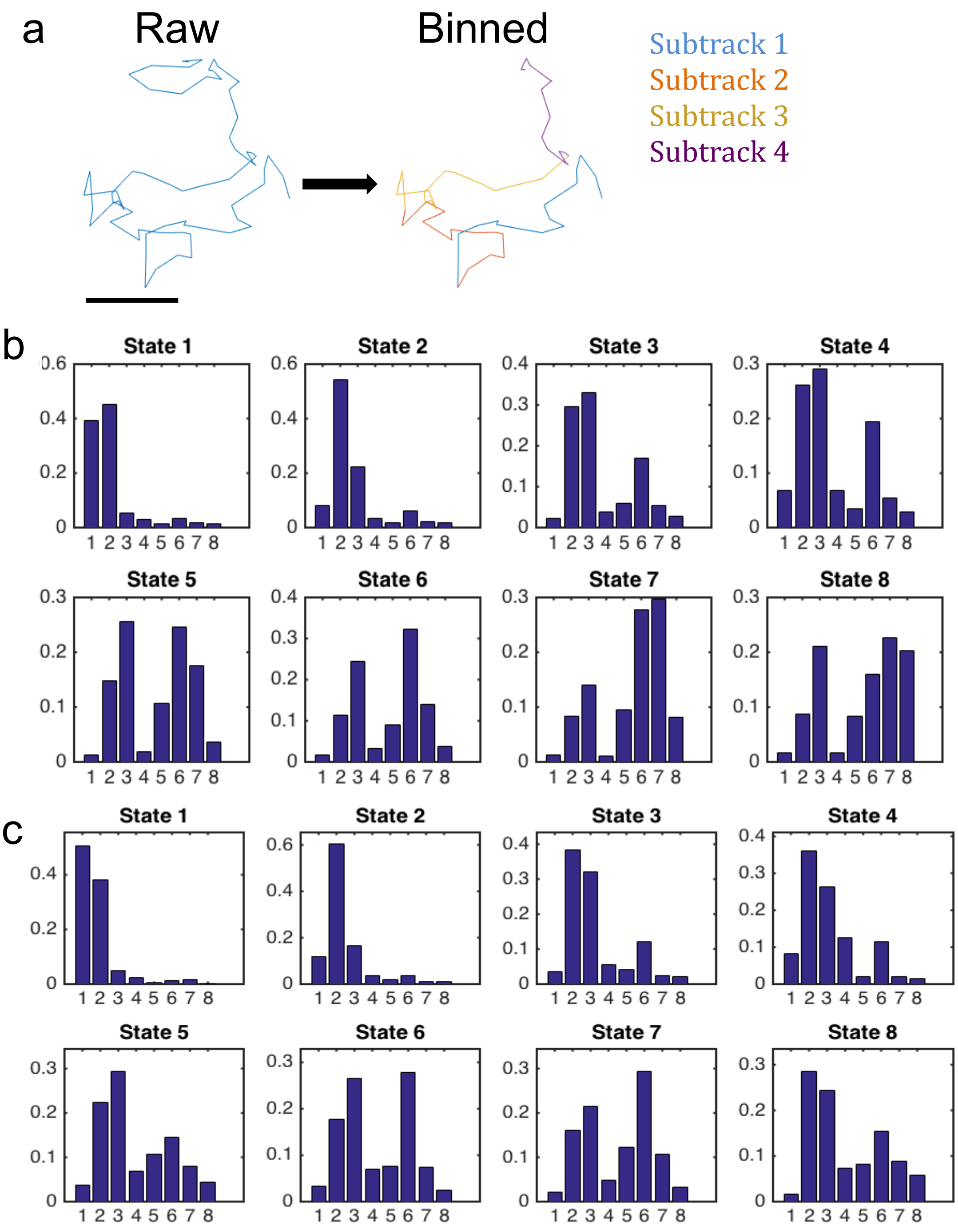


**Supplementary Figure S3** a) All single BCR molecule trajectories are split into 15 frame long sub-tracks. pEM analysis is performed over the set of 15 frame long binned tracks and then the original trajectories are reconstructed to obtain information about the transitions of molecules across different states. Scale bar is 1 μm. b) Plots showing the fraction of transitions made from each state to itself and to all others in control cells. Numbers on the x-axis indicate the state to which the transitions are being made. c) Plots showing the fraction of transitions made from each state to itself and to all others in cNKO cells.

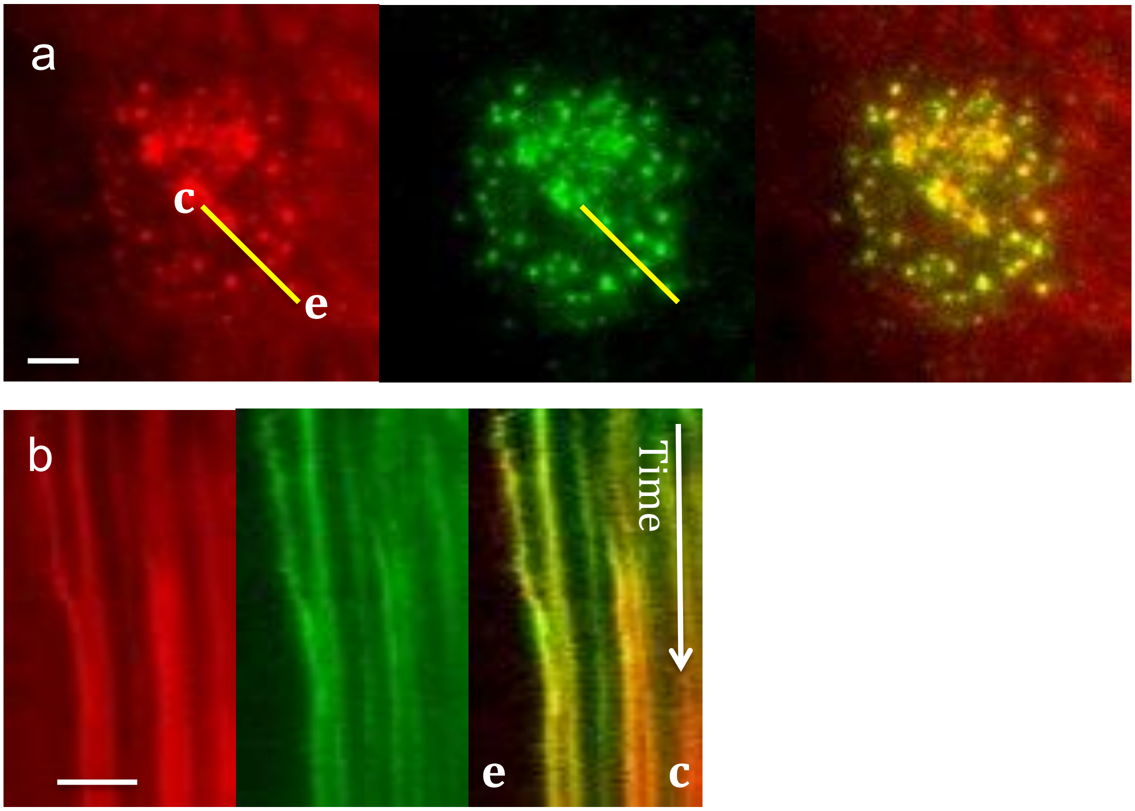


**Supplementary Figure S4.** a) TIRF images of BCR (red) and CD19 (green) microclusters and the overlay of both channels. b) Kymograph generated along the line indicated in a) showing the inward movement of the microclusters and their superposition as they move. **c** marks the center of cell and **e** the cell edge.

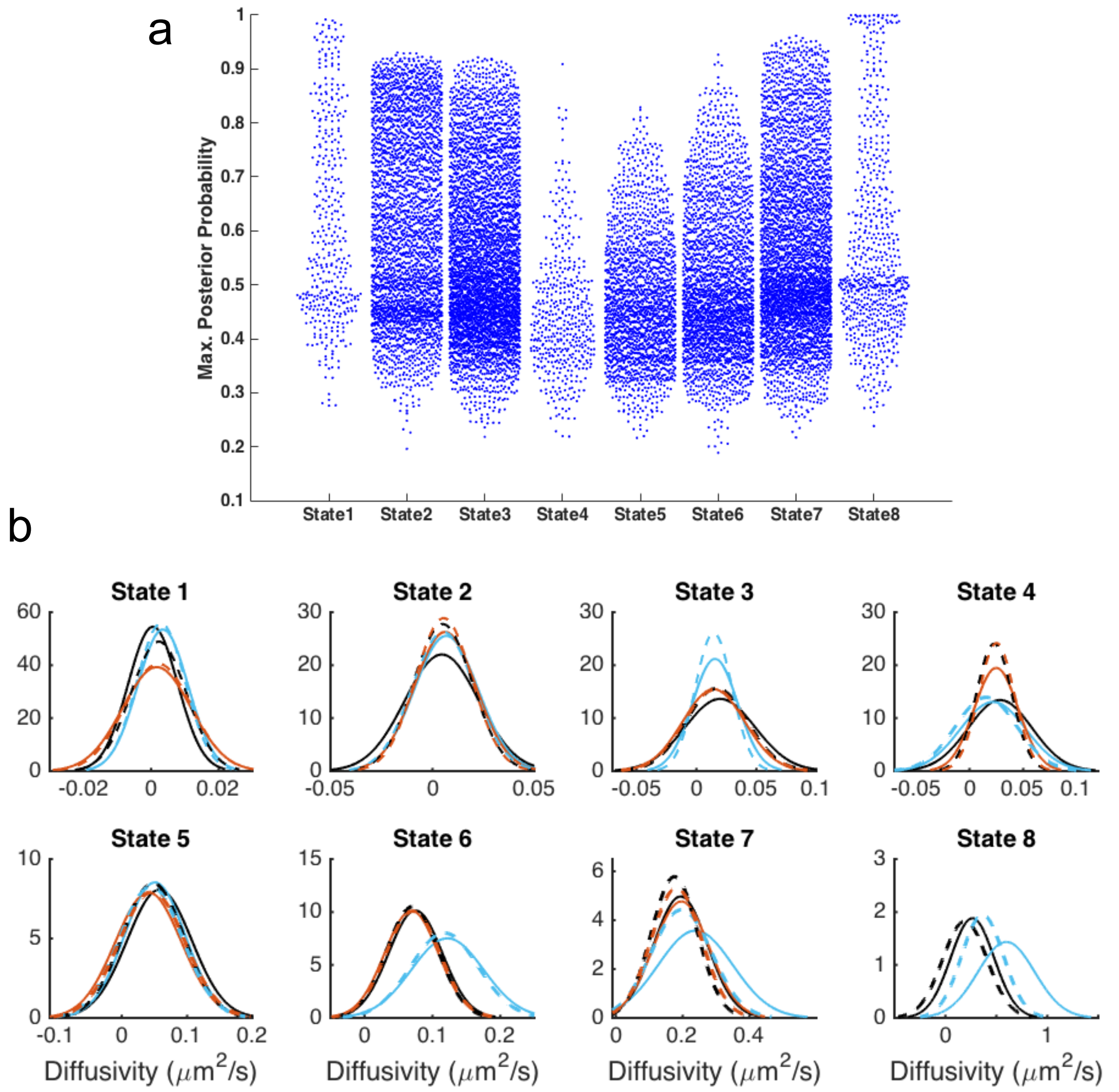


**Supplementary Figure S5** a) Beeswarm plots showing the maximum posterior probability used to assign a track to a particular state for BCR in control cells. Each point represents a 15 frame long track. b) Plots of diffusivity distributions for each state used to compare across conditions. Black curves correspond to BCR, blue to CD19 and orange to FcγRIIB. The solid lines correspond to control cells and the dashed lines to cNKO cells.
